# Pharmacogenomic discovery of genetically targeted cancer therapies optimized against clinical outcomes

**DOI:** 10.1101/2024.01.05.574245

**Authors:** Peter Truesdell, Jessica Chang, Doris Coto Villa, Meiou Dai, Yulei Zhao, Robin McIlwain, Stephanie Young, Shawna Hiley, Andrew W. Craig, Tomas Babak

## Abstract

Despite the clinical success of dozens of genetically targeted cancer therapies, the vast majority of patients with tumors caused by loss-of-function (LoF) mutations do not have access to these treatments. This is primarily due to the challenge of developing a drug that treats a disease caused by the absence of a protein target. The success of PARP inhibitors has solidified synthetic lethality (SL) as a means to overcome this obstacle. Recent mapping of SL networks using pooled CRISPR-Cas9 screens is a promising approach for expanding this concept to treating cancers driven by additional LoF drivers. In practice, however, translating signals from cell lines, where these screens are typically conducted, to patient outcomes remains a challenge. We developed a pharmacogemic (PGx) approach called “Clinically Optimized Driver Associated PGx” (CODA-PGx) that accurately predicts genetically targeted therapies with clinical-stage efficacy in specific LoF driver contexts. Using approved targeted therapies and cancer drugs with available real-world evidence and molecular data from hundreds of patients, we discovered and optimized the key screening principles predictive of efficacy and overall patient survival. In addition to establishing basic technical conventions, such as drug concentration and screening kinetics, we found that replicating the driver perturbation in the right context, as well as selecting patients where those drivers are genuine founder mutations, were key to accurate translation. We used CODA-PGX to screen a diverse collection of clinical stage drugs and report dozens of novel LoF genetically targeted opportunities; many validated in xenografts and by real-world evidence. Notable examples include treating STAG2-mutant tumors with Carboplatin, SMARCB1-mutant tumors with Oxaliplatin, and TP53BP1-mutant tumors with Etoposide or Bleomycin.

**One Sentence Summary:** We identified principles of pharmacogenomic screening that predict clinical efficacy in cancer patients with specific driver mutations.

## INTRODUCTION

One in six deaths in the United States is caused by cancer. Yet, despite enormous investments of time and money, survival rates for high fatality cancers have not significantly improved in the past 50 years (*1*). This is caused in part by the high development costs for new therapies and low success rates of clinical trials. In fact, despite massive advances in technology and a 100x increase in spending, more than 90% of drugs entering clinic trials do not reach the market (*2*). Importantly, most candidates do not fail because of drug toxicity, but because of a lack of efficacy; this results in thousands of potential treatments being shelved late in the clinical trial process, including more than 2,400 cancer therapeutics that were shelved after being tested in phase 2 clinical trials (**Supplemental Figure 1** and see ClinicalTrials.gov (*3*)).

Standard chemotherapy is one of three historically dominant approaches to cancer treatment, which continue to be frontline therapies for most patients today, along with surgery and radiation (*4*). Immunotherapy and targeted therapies are more recent approaches that have yielded improved outcomes for some patients. As cancer is a collection of more than one hundred distinct subtypes (*5*), treatment has benefited from extensive efforts to catalog the molecular subtypes and causal genetic mutations (*6*). Targeted therapies are designed to exploit cancer dependencies and are based on the assumption that the molecular features of a patient’s tumor influence clinical response. This targeted approach to cancer therapy may lead to increased efficacy and reduced toxicity compared to chemotherapy treatments, which do not selectively target cancer cells and thus have considerable off-target effects. Indeed, the notion that drugging the causal biology is effective is reflected in the clinical success of targeted therapies: more than 50 are currently approved, yielding more than $100 billion in global sales, projected to grow to $258 billion by 2032 (*7*).

Tumor genomics studies have identified cancers linked to specific driver mutations that confer selective advantages to cancer cells. Driver mutations are highly predictive of drug activity across multiple cancer types. Evidence for the effectiveness of this approach is that most accelerated approvals for treatments over the past 20 years have been targeted therapies (*6*), some of which have yielded overall response rates of >80% (e.g., Selpercatinib in lung cancers with RET mutations (*8*)). The efficacy of these targeted treatments is attributed to the fact that they inhibit the causal biology of cancers.

Approximately one third of cancers are caused by driver mutations that induce a gain of function (GoF) in oncogenes and result in a protein product that can be therapeutically targeted. The remaining two thirds of cancers are caused by a loss of function (LoF) mutation in a tumor suppressor gene (TSG). In healthy cells, TSGs control cell growth and division; LoF mutations in these genes result in uncontrolled growth and cancer (*9*). Such cancers are much harder to target because it is the absence of a functional protein that results in oncogenesis. As evidence of this disparity in treatability, approved drugs exist for treating ~60% of cancers caused by oncogenic activation, but only ~2% of TSG-driven cancers (**Supplemental Figure 2**). One approach to targeting cancers driven by LoF mutations is to leverage their unique vulnerabilities by identifying synthetic lethal (SL) interactions in which the relationship between two genes with LoF in either, but not both is tolerated. Since only tumor cells contain the driver mutation, drugs that specifically target such a mutation will impact only these cells, and toxicities are expected to be minimal. This approach has been effectively applied to identify both genetic dependencies and small molecule sensitivities of cancer cell lines (*10*). As evidence of the importance of this approach, massive and ongoing efforts by large consortia are underway to comprehensively map SL networks in hundreds of cancer cell lines with the goal of identifying drug targets and the patients who are expected to respond best to targeted treatment (*11*).

While these efforts have provided valuable insights and identified potential drug candidates, significant challenges remain. Most patients still do not benefit from targeted therapies because: (i) large-scale screens produce new target predictions and feed drug development pipelines at the entry point such that treatment is years away, and (ii) genetic SL relationships identified in cell lines may not translate to tumors (*10*). A recent analysis revealed that SL connections identified from large-scale pooled CRISPR screens are more likely to replicate SL potential in tumors when a driver gene is involved (*12*). It has also been demonstrated that pharmacologic inhibition of a cell line rarely reproduces the phenotype induced by genetic knockout (KO) of the drug target (*12, 13*). This suggests that it is not possible to identify drugs with potential to be repurposed as genetically targeted therapies by simply aligning drugs against novel targets discovered on the basis of genetically-defined SL. However, pairing these approaches with pharmacogenomic screens has shown promise to define druggable targets that are essential with a particular LoF TSG (*14, 15*).

Here we describe CODA-PGx (clinically optimized driver associated PGx), a pharmacogenomic platform developed to accurately predict utility of small molecule drugs as genetically targeted therapies. CODA-PGx simultaneously tests drug viability interactions against all common driver genes generated as a pool of isogenically controlled cell lines. Screening drugs with extensive clinical outcome data enabled us to discover and optimize parameters that are relevant to predicting clinical-stage efficacy. We show that modeling driver biology in cell lines is key to accurate prediction of genetic sensitizers relevant to clinical treatment. This also applied to patient selection, where the therapy is likely to work best if targeting a foundational driver mutation that is causal to tumor initiation and progression. We expect the principles described in this study to aid in the discovery of therapeutic opportunities that emerge from pharmacogenomic screening.

## RESULTS

### Pooled CRISPR pharmacogenomic screening to identify druggable driver-mutation dependencies

Targeted therapies are designed to treat tumors with driver mutations in either oncogenes (GoF) or tumor suppressor genes (LoF). Unlike oncogenes, where mutations result in a targetable protein, LoF mutations in tumor suppressor genes typically yield reduced functionality of the protein that cannot be easily reversed pharmacologically (**Figure 1A**). Consequently, the majority of approved genetically targeted therapies are for cancers driven by oncogene mutations (**Supplemental Figure 1**); whereas there remains a major unmet need for treatment of tumors driven by mutation of tumor suppressor genes.

**Figure 1:**
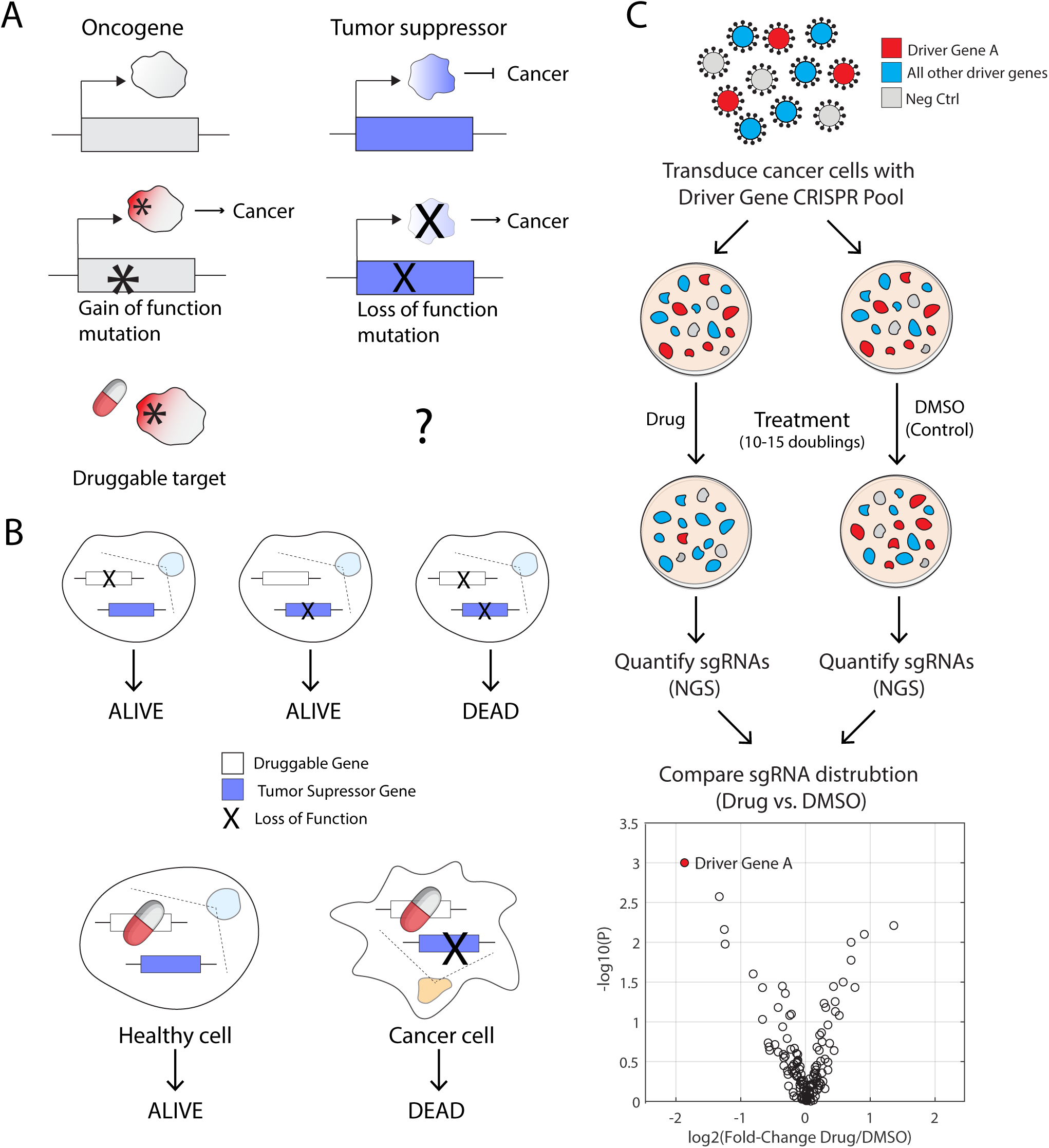
CRISPR-pooled pharmacogenomic screening to identify druggable genetic dependencies of LoF TSG mutations. **(A)** Schematic showing the drug-targetability of GoF versus LoF mutations. GoF mutations result in a protein product that can be pharmacologically targeted, while LoF mutations results in the absence of a protein product. **(B)** Schematic showing the synergistic effects of SL mutations on cell viability. Single mutations in either green or blue genes do not impact cell growth, but when both are mutated in the same cell the combination is lethal. (**C)** Schematic of CODA-PGX platform for rapid, high throughput, pharmacogenomic screening. Cancer cell lines are transduced with a pool of driver gene sgRNAs to create knockouts via CRISPR and grown in the presence or absence of drug. The viability of each knockout is detected by NGS sequencing of the corresponding guide RNAs used to create it. Data is visualized by comparing sgRNA distribution between drug and control samples.

It is widely accepted that identifying vulnerabilities of tumors through SL interactions is key to unlocking the potential of TSG targeting (**Figure 1B**) and driver genes have recently become recognized as important to clinical translation (*12*). While large consortia have made great advances to identify genes which when knocked out together kill cancer cells, challenges remain. Despite the availability of whole-genome pooled CRISPR screening data from more than 1000 cell lines, we are likely severely underpowered to complete a comprehensive mapping of SL networks without the use of isogenic screens (*16*). Furthermore, translating SL discoveries in cell lines is challenging for both scientific (*12*) and practical (*2*) reasons.

### CODA-PGX platform design

To address these challenges, we developed a method that efficiently identifies sensitizing driver mutations for small molecule drugs (**Figure 1C**). In brief, the approach is to: (i) perform driver-focused CRISPR screens with and without drug in a pool of isogenically controlled cell lines, (ii) use NGS to identify genes which, when mutated, cause cells to ‘drop out’ of the pool. We refer to the genes containing these mutations as ‘hits’ and expect that their presence in tumors may identify patients whose clinical outcomes could be improved by treatment with the corresponding drug. We curated driver genes represented on our platform (**Supplemental Table 1**) from published lists (*17–19*) and subset to genes mutated in at least 5% of one cancer type in The Cancer Genome Atlas (TCGA) (*20*). 95.1% of all patients in TCGA have at least one mutation in a gene targeted on our platform. 3-4 guide sequences targeting these genes were selected from the Brunello library (*21*). Importantly, we also added 96 guides that target non-conserved intergenic regions to serve as negative controls (**Supplemental Table 2**).

To visualize the data, we plot, for each driver gene, the mean ratio of guide sequences present in drug-treated cells versus guide sequences present in control (vehicle-only) on a log scale (**Figure 1C, bottom**). Guide sequences with negative log values represent genes which when mutated reduce the viability of the cells in the presence of the drug. We then gauge significance (y-axis) using a ranksum test of the 3-4 guides/gene ratios against the distribution of ratios of the 96 negative controls. Negative controls establish a conservative null distribution of platform technical variation, thereby yielding a robust drug-driver synergy signal.

Motivated by the recent discovery that driver biology is important for translating discoveries made in cell lines to clinical outcomes (*12*), we optimized several parameters to maximize representation of native driver biology. We found that oncogene GoF and TSG LoF effects on viability are reproduced in public genome-wide pooled CRISPR screens such as the Dependency Map ((*22, 23*); **Supplemental Figure 3**). For example, cell lines with an oncogenic BRAF^V600E^ activating mutation were more susceptible to BRAF-knockout (KO) than BRAF-wildtype (WT) lines, and KO of the TSG PTEN in PTEN-WT lines increased the relative fitness of cells. We therefore identified, for each driver gene, the set of cell lines where these expected effects on proliferation were maintained. We also used Celligner (*24*) to identify cell lines whose expression profile matches its tumor of origin. Furthermore, to best capture the founder-driver biology of our genetic perturbations, we reasoned that TSGs present as 2 WT copies, with evidence of expression (*25*) best represent the native initial tumor state.

In summary, a “valid” TSG perturbation in our platform: 1) results in a fitness increase when knocked out in DepMap, 2) is expressed from two WT copies, and 3) is in a cellular context that matches its tumor of origin. Since pooled CRISPR screens are not yet empowered to generate specific edits, we rely on the GoF mutation to be present in the cell line background. For a GoF oncogene to be “valid” signal in our CODA-PGX platform, it similarly needs to yield a fitness effect in DepMap; this time negative demonstrating the dependency on that oncogene. It similarly needs to be expressed (*25*), present in 2 or more copies, and the cell line needs to reflect the tumor of origin. We then optimized cell line selection to maximize representation of “valid” perturbations (see **Methods** for details) and quantified screening results on “valid” perturbations whenever possible.

### CODA-PGX platform testing and validation

To test if our approach could accurately identify known SL interactions, we performed proof-of-concept experiments with four currently marketed PARP (poly-ADP ribose polymerase) inhibitor drugs - the only approved class of drug currently available to treat LoF TSG drivers. PARPs are nuclear proteins that detect and initiate a cellular response to repair single strand breaks in DNA, and PARP inhibitors cause cells to accumulate DNA damage and results in cell death. Accordingly, PARP inhibitors have established SL interactions with genes involved in homologous recombination, such as BRCA1, BRCA2, ATM, ATR, RAD21, and STAG2 (*26, 27*).

In order to validate the platform, establish parameters for selecting an appropriate drug screening concentration, and to explore the effect of cell population doubling, we screened all four currently approved PARP inhibitors in several cell line backgrounds (see Methods) while varying the above parameters. We examined viability inhibitory concentration (IC) values from IC_5_-IC_50_ and population doublings from 5-15. We determined that IC_15_ and 7-10 population doublings are optimal to maximize the signal of known clinically validated SL interactions (**Supplemental Figure 4**; **Figure 2**). At lower concentrations there are no significant signals, and at higher concentrations general drug toxicity/resistance effects start to dominate. For example, for hits above IC_30_, metabolic genes (e.g. mTOR) and genes involved in more general biology (e.g. DIS3, an exoribonuclease), reproducibly dominate the most significant sensitizers (**Supplemental Figure 4**).

**Figure 2:**
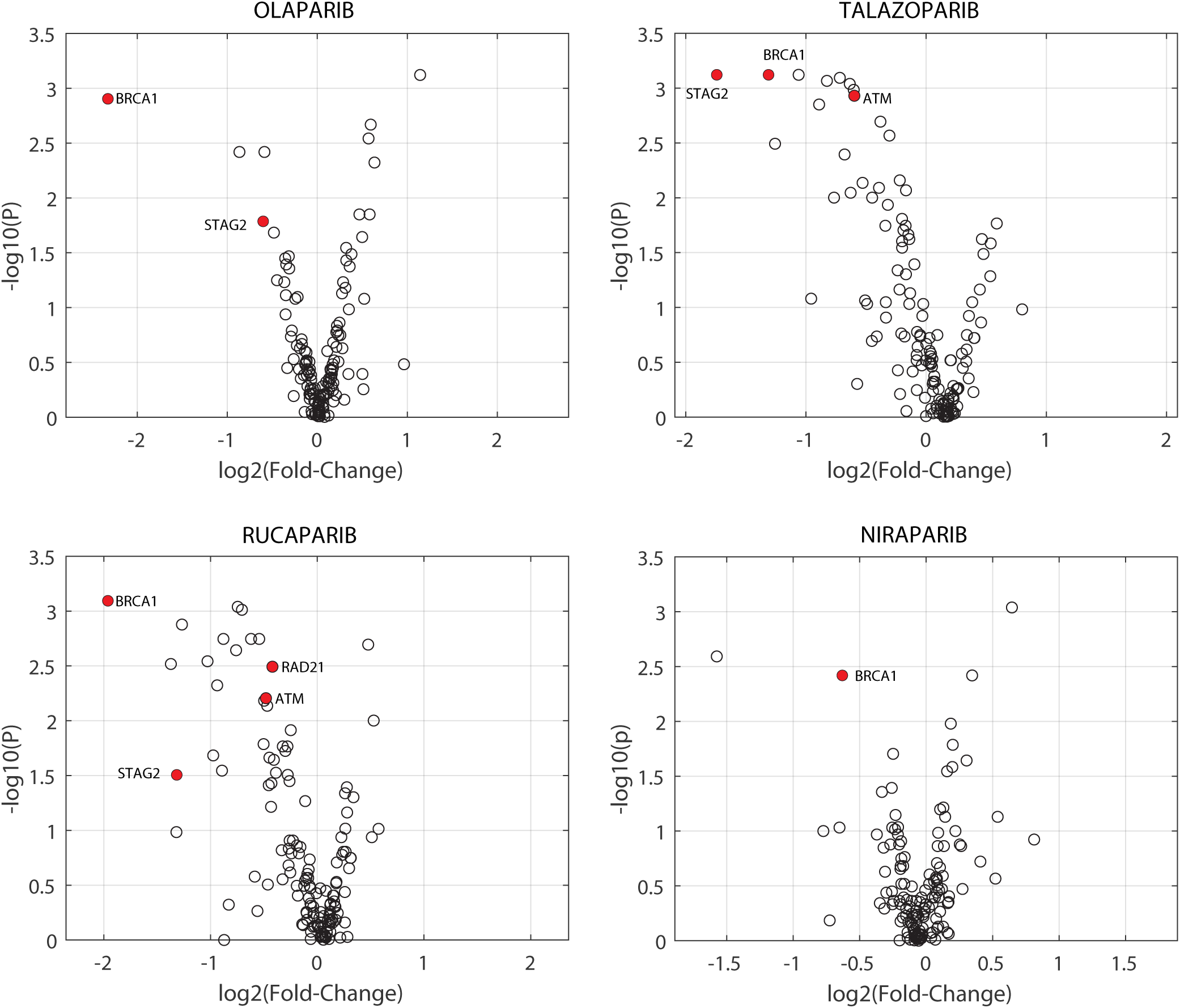
CODA-PGX platform proof-of-concept experiment with PARP inhibitorsin MDA-MB-231 cells. Volcano plots (see **Methods**) depict SL interactions between driver mutations and four currently marketed PARP inhibitor pharmaceuticals. Known SL interactions that were identified by CODA-PGX are highlighted in red.

### Mutation-induced drug sensitivity landscape of common chemotherapeutics

To discover novel patient selection possibilities for existing treatments, we screened a selection of marketed and characterized drugs. Specifically, we analyzed SL interactions of our pooled driver mutations in 85 cell line-drug combinations and visualized the results in a heatmap (**Figure 3A, Supplemental Table 3**) where cell line-drug combinations are plotted on the y-axis and driver mutations on the x-axis. Yellow squares indicate sensitization in the presence of drug (i.e., slower growth compared to control), while blue squares indicate that a drug conferred a fitness benefit (i.e., faster growth compared to control). Data were clustered and diagonalized to highlight specific drug-driver combinations along the diagonal (*28*). Vertical stretches of yellow squares indicate that a driver mutation was significantly sensitizing in multiple screens. In general, these patterns correspond to drugs within a single class (e.g., PARP inhibitors: Olaparib, Rucaparib and Talazoparib; **Figure 3A, Box 1**).

**Figure 3:**
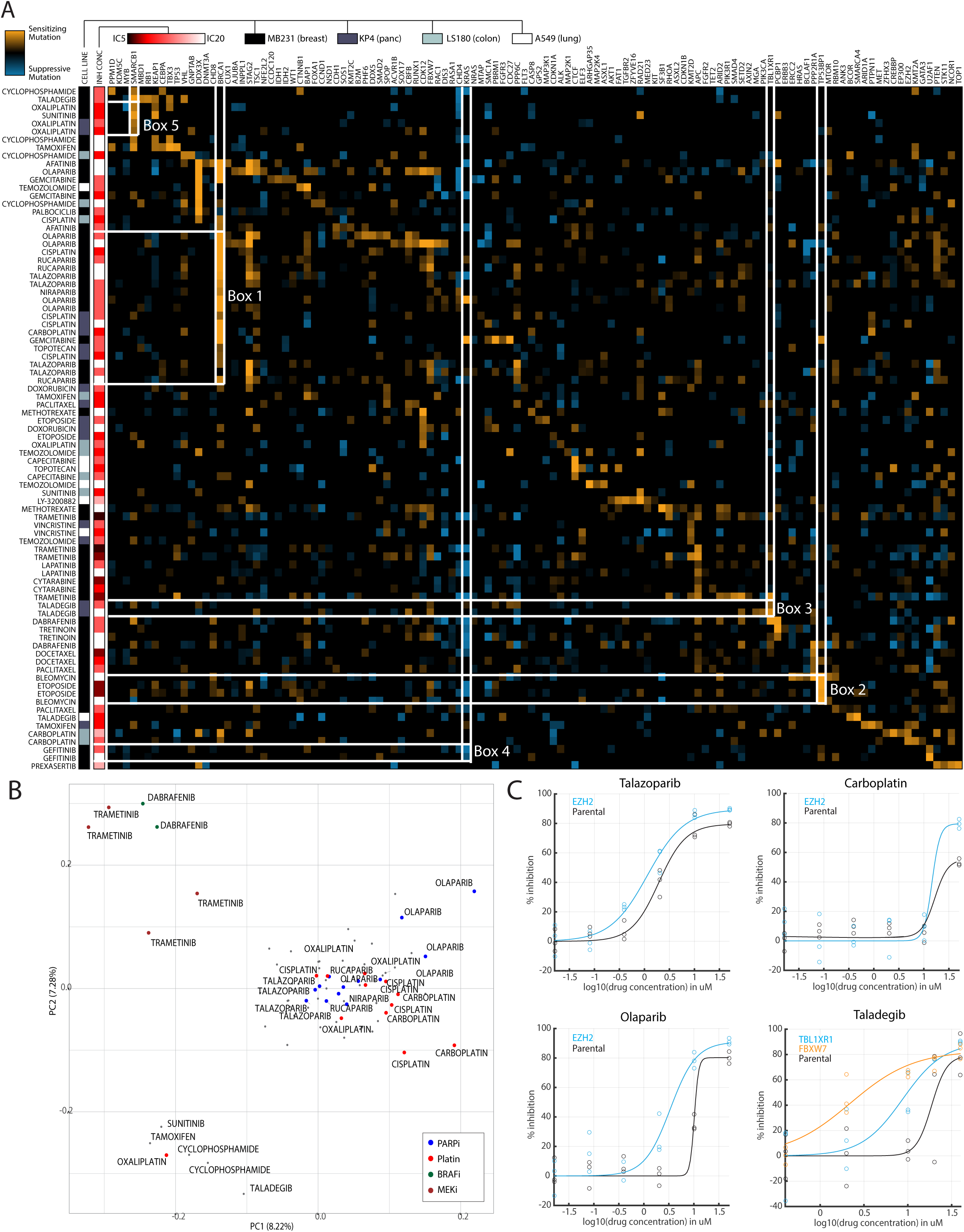
Mutation-induced drug sensitivity landscape of common chemotherapeutics. **(A)** Heatmap of driver mutations (x axis) and cell line/drug combinations (y axis) where yellow squares indicate reduced fitness in the presence of drug and blue squares indicate increased fitness. **(B)** PCA of interactions identified by CODA-PGX. **(C)** Dose response curves of selected PARPi and platinum-based chemotherapy drugs with and without sensitizing mutations identified by CODA-PGX.

Interestingly, we note that both Cisplatin and Carboplatin also occur within this same vertical pattern, suggesting that they have similar SL interactions to PARP inhibitors. Platinum-based chemotherapies are a class of chemotherapy drugs whose mechanism of action involves delivery of platinum ions to cells that cause DNA crosslinks which inhibit DNA replication and synthesis in all dividing cells. Although not currently prescribed as targeted therapy, platinum-based chemotherapies have been associated with improved outcomes in BRCA1- and BRCA2-deficient tumors (*29–31*).

A second group of SL interactions is detected between the TP53BP1 gene and etoposide and bleomycin (**Figure 3A, Box 2**). TP53BP1(tumor-suppressor p53 binding protein 1) has a known role in promoting the non-homologous end joining pathway to repair DNA double strand breaks (*32, 33*). Etoposide and bleomycin are both chemotherapy drugs, often used in combination, known to impact DNA damage and repair; etoposide prevents the repair of DNA damage while bleomycin causes DNA degradation. While neither drug is currently used in targeted therapy approaches, the observed SL interaction between these drugs and TP53BP1 is consistent with reports in the literature (*34–36*) and with the mechanism of action of the drugs and biological role of the TP53BP1 gene.

We also noted a SL interaction between TBL1XR1 and Taladegib (**Figure 3A, Box 3**). Taladegib is a drug that was developed to inhibit Hedgehog (Hh) signaling, a pathway often upregulated in cancer cells (*37*). Specifically, Taladegib is a small molecule antagonist to the smoothened receptor (an upstream component of Hh signaling that activates pathway members to promote cell division, differentiation, and migration). It has been shown to inhibit proliferation of tumor cells where the Hh pathway is abnormally active. TBL1XR1 has been identified as a potential therapeutic target (*38*) as mutations are widely documented in cancer (*39*). While we are not aware of a direct connection between TBL1XR1 and Hh signaling, the gene is tightly linked to Wnt signaling (*40*) and there is well-established crosstalk between these two signaling pathways. Further, Hh signaling is highly activated in B cell lymphoma (*41*), which is also driven by TBL1XR1 mutations (*39*). The crosstalk between Wnt signalling and Hh signaling is well documented, and the two pathways often perform similar biological functions (i.e., are highly influential for cell division, differentiation, and migration). TBL1XR1 function is important for proper Wnt signaling, so its KO (to disrupt Wnt signaling) combined with Taladegib treatment (to inhibit Hh signaling) is expected to be SL for cancer cells.

These data also allow us to make some general observations about the value and potential impact of CODA-PGX. We note that many of the driver mutations identified as SL hits are consistent with the known mechanism of action of the drugs tested. Specifically, both platinum-based chemotherapy and PARP inhibitors are SL with chromatin-associated proteins STAG2 and EZH2, and BRCA1 (known to function in DNA damage repair). We observe smaller but significant interactions between Taladegib and FBXW7 and SMARCB1 genes, both of which are known to regulate Hh signaling (*42, 43*). Additional noteworthy interactions are summarized in **Supplemental Table 4**, including a variety of suppressive interactions. For example, MDA-MB-231 cells, which have an activating KRAS mutation, were less sensitive to Gefitinib (**Figure 3A, Box 4**), a previously reported effect observed in the clinic (*44*). Taken together, these observations suggest that in addition to identifying established SL interactions between drugs and driver genes, CODA-PGX uncovers new and biologically relevant connections.

To explore whether drug sensitizing mutation signals arising from CODA-PGX screening are confounded by any technical variables, we performed a PCA analysis (**Figure 3B**, see Methods). Briefly, the proximity of data points (each of which represents one screen – or cell line/drug combination) in the plot reflects how similar they are. We observe that, in general, drug mechanism of action is the dominant variable that drives the signal, not the cell line in which the screen was conducted, or the drug concentration. For example, consistent with the heatmap (**Figure 3A**), we observed that PARP inhibitors and platinum-based chemotherapy cluster tightly together, as well as clustering of drugs with other unique mechanisms (**Figure 3B**). This suggests that the druggable dependencies arising from cancer driver biology, when filtered appropriately (e.g. on driver context; see Methods), tend to persist across different cell of origin contexts. To orthogonally validate some novel hits from our CODA-PGX screens, we generated isogenic pairs of cell lines differing only by the CRISPR/Cas9-mediated KO of the indicated driver gene in MDA-MB-231 cells (EZH2) or KP4 cells (TBL1XR1, FBXW7). We then compared parental and KO isogenic lines for their sensitivity to drugs tested in our CODA-PGX platform. In all cases, we observed greater inhibition of growth in the KO lines compared to parental lines at the same drug concentration (**Figure 3C**). This confirms that sensitizing driver mutations identified by our CODA-PGX platform indeed reduce cellular fitness in the presence of drug.

### In vivo validation of STAG2 – a novel Carboplatin-sensitizing mutation

We next investigated the ability of CODA-PGX to identify driver mutations that improve the efficacy of specific drugs. We looked specifically at Carboplatin, a chemotherapy drug commonly used to treat ovarian and other cancers, but not designated for targeted therapy. Our preliminary screening and heatmap analysis (**Figure 3**) identified that Carboplatin shares several SL interactions with PARP inhibitor drugs. To explore whether there are additional driver mutations that improve the efficacy of Carboplatin, we screened for SL interactions in three different cell lines using our CODA-PGX approach (**Figure 4A and Supplemental Figure 5**). In addition to BRCA1, mutations in STAG2, ATM, DDX3X, EZH2, and APC showed decreased viability in the presence of carboplatin compared to the vehicle control (**Figure 4A**). Of these, ATM, STAG2 and EZH2 have known involvement in homologous recombination and DNA repair (*26, 27, 45*). Since STAG2 was identified as a strong hit and is not currently used as an HRD patient-selection marker, we pursued more extensive validation of its potential as a sensitizing mutation. We also noted from our data that the KO of TP53 causes resistence to Carboplatin (**Figure 4A**) and aligns with documented resistance mechanisms in the clinic (*46*).

**Figure 4:**
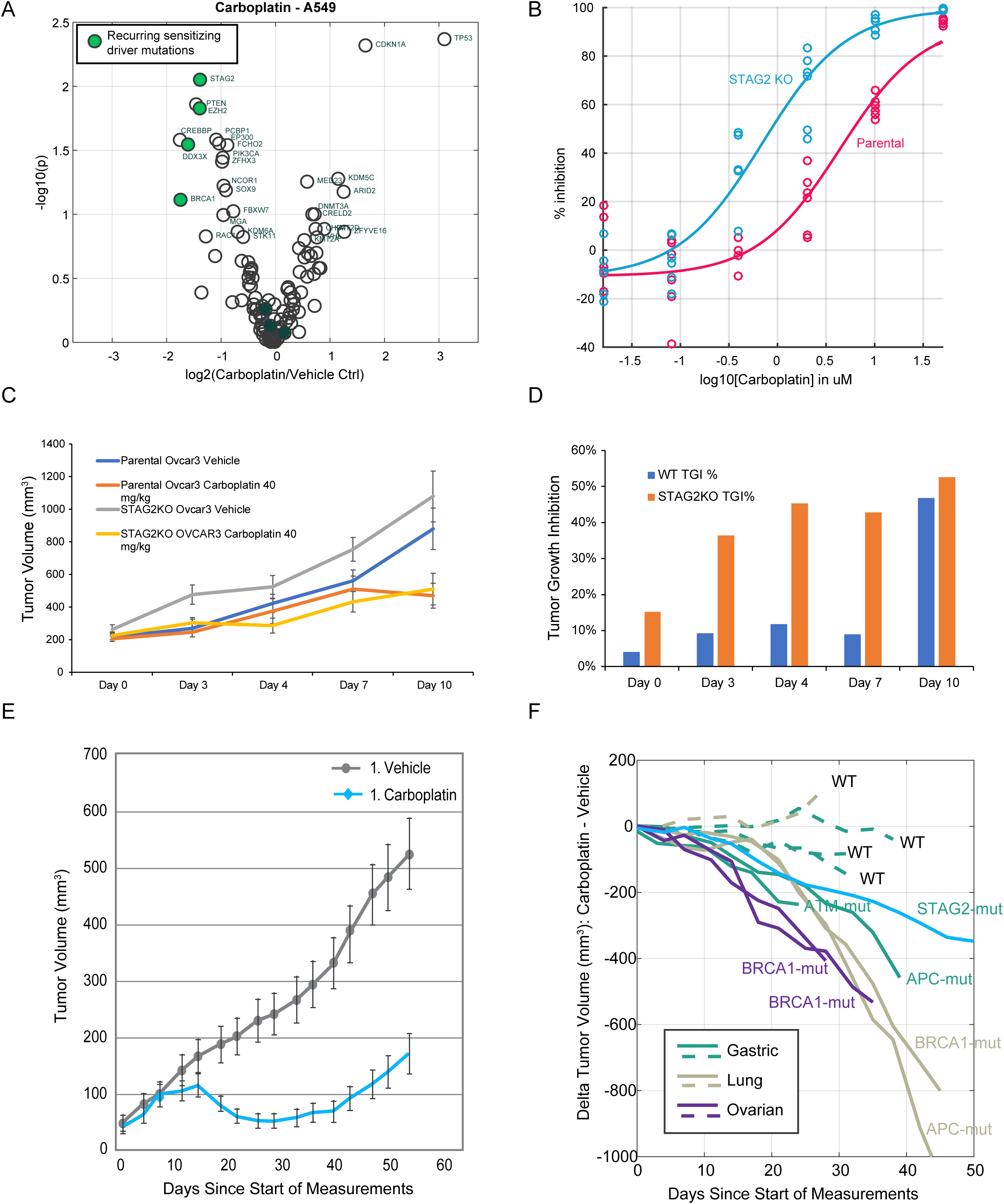
In vivo validation of STAG2 – a novel Carboplatin-sensitizing mutation. **(A)** Volcano plot identifying sensitizing mutations detected in A549 cells. **(B)** Dose response curve showing inhibitory effects of Carboplatin on STAG2 KO (blue) and Parental (pink) OvCar-3 cells. **(C)** Tumor growth curves of cell-derived xenografts (CDXs) from STAG2 KO and Parental OvCar-3 cells with and without Carboplatin treatment. n=10 mice per group, data presented as the mean±SEM. A two-way ANOVA test was used. Parental OvCar-3 Vehicle v.s. Carboplatin, *p* = 0.17; STAG2 KO Vehicle v.s. Carboplatin, *p* = 0.0056; \**P* < 0.05 was considered statistically significant. **(D)** Normalized tumor growth inhibition (TGI) values for WT and STAG2 KO CDXs shown in (C). **(E)** Plot of tumor volume over time for a gastric STAG2 LOF model (CrownBio model GA0151) patient-derived xenograft (PDX). n= 8 mice per group. **(F)** Plot of tumor volume over time for additional Crown Bio PDX models of gastric, lung, and ovarian cancer cells. Turquoise line coresponds to model shown in (E).

We first screened STAG2 LoF in a controlled viability assay. We compared the growth of cells with a CRISPR-generated KO of STAG2 to parental cells in the OvCar-3 ovarian cancer cell line (see **Methods**) in presence of increasing concentrations of Carboplatin and plotted a dose-response curve (**Figure 4B**). The results confirm that the absence of STAG2 causes greater growth inhibition at lower concentrations of Carboplatin, indicating a SL response between STAG2 and Carboplatin.

Cancer cell-derived xenografts (CDX) provide a valuable system in which to validate the genetic sensitization of LoF mutations in vivo. We established CDXs in mice by injecting NSG mice with either WT or STAG2 KO OvCar-3 cells. We observed that the CRISPR-generated STAG2 KO cells grew faster than WT cells (**Figure 4C**, KO vehicle vs WT vehicle). However, the observed inhibitory effect within the same genetic background indicate that Carboplatin has a greater inhibitory effect on the growth of STAG2 KO tumors. We calculated the normalized effect of Carboplatin on STAG2 KO vs WT as tumor growth inhibition and plotted these data (**Figure 4D**). We observed that the normalized effect of Carboplatin on STAG2 mutant growth is greater than the normalized effect of Carboplatin on WT cells. The study was ended on Day 10 because of the fast growth rates of this tumor model; we speculate that the growth slowed as tumors reached a maximum size, and that this may explain the reduction in tumor growth inhibition (TGI) at this timepoint (**Figure 4D, Day 10**).

We next used a patient-derived xenograft (PDX) system to further validate these observations in a pre-clinical system that closely models clinical response. PDX models generally capture more of the heterogeneity of tumors compared to CDX models and have been shown to have reasonable predictive capacity for genetically-guided precision oncology (*47*). To this end, we partnered with Crown Bio to interrogate their collection of Carboplatin standard-of-care in ovarian, gastric, and lung cancer PDX models. We plotted tumor volumes from model GA0151, a gastric tumor PDX with STAG2 LoF mutant model treated with vehicle or 40 mg/kg Carboplatin Q4Dx4 (**Figure 4E**) and observed that the presence of the LoF mutation slowed tumor growth compared to WT when treated with Carboplatin. We expanded this analysis to examine tumors in the ten available models for these three cancer types and plotted the difference in tumor volume between WT and mutant for BRCA1, ATM, STAG2, and APC mutants (**Figure 4F**). In all cases, the presence of the driver mutation in the PDX showed reduced tumor growth with Carboplatin compared to WT PDXs of the same cancer type. These data indicate that the mutations identified by CODA-PGX were sensitive to Carboplatin and, by extension, that CODA-PGX can accurately identify sensitizing mutations.

### Retrospective clinical validation of hits

The ultimate test for a new therapy is clinical efficacy, and the gold standard measure of this is a prospective randomized control clinical trial. In the last 10 years we have seen a dramatic increase in cancer patient data collection, including NGS/molecular readouts that capture the mutation spectrum of the patient’s tumor, treatment, and outcomes. The Cancer Genome Atlas (*48*) contains more than ten thousand patients with tumors profiled by whole-exome sequencing and their treatment histories and outcome meticulously documented. More recently, the AACR’s Project Genie is a cancer registry of patient genomic and demographic data for more than 150,000 patients across 19 oncology centers (*49*). While molecular profiling in Project Genie is largely limited to targeted sequencing of cancer associate genes, and drug treatment/outcome data is not yet broadly available, an industry-academia partnered effort known as the Biopharma Collaborative (BPC) (*50*) is gradually releasing these curated RWE datasets. We downloaded all mutation, treatment history, and clinical outcome data from TCGA for 33 cancer types and BPC data from lung and colon cancers that were available at time of publication.

To evaluate the ability of CODA-PGX to predict drug-driver interactions likely to yield meaningful improvements in clinical outcomes, we considered the input data from our exploration of the mutation-induced drug sensitivity landscape of common chemotherapeutics (**Figure 3**). For each cell line/drug combination (i.e., row), we identified genes that sensitized the cells to drug treatment (negative on volcano plot) at p<0.05 and a minimum fold-change of 2. In order to avoid potentially confounding variables arising from cross-cancer analysis, we selected the cancer type with the highest number of patients where the drug in question was used as a frontline therapy. We then subset these patients into two groups based on their tumor mutation status: with or without one or more of the sensitizing mutations identified in the screen (see Methods for details). We then compared overall survival (OS) of these populations based on driver mutation status (**Figure 5A, B**). With our CODA-PGX hit linking SMARCB1 LoF to Oxaliplatin efficacy (**Figure 3A**, **Box 5**), we compared overall survival of colorectal cancer patients who had mutations to the SMARCB1 gene and observed that patients with mutations treated with Oxaliplatin (**Figure 3A**, **Box 5**) survived longer than patients without SMARCB1 mutations (**Figure 5A**, p=0.033, logrank test). A parallel analysis of glioma and glioblastoma patients treated with Temozolomide revealed significantly improved survival in patients with CODA-PGX-informed driver mutation-positive tumors compared to patients without the indicated driver mutations (**Figure 5B**, p<0.001, logrank test). We also repeated the above analysis for mutations that suppressed drug sensitivity (positive in volcano plots, blue squares in **Figure 3**; e.g. **Figure 5C**). We observed significantly decreased probability of survival in non-small cell lung cancer patients with these mutations compared to patients without the mutations who received the same treatment (p=0.05, logrank test). To confirm that the mutations themselves are not responsible for the difference in OS, whenever possible (i.e., data exists), we repeated this analysis for the next best represented drug and observed no difference in overall survival (**Supplemental Figure 6**).

**Figure 5:**
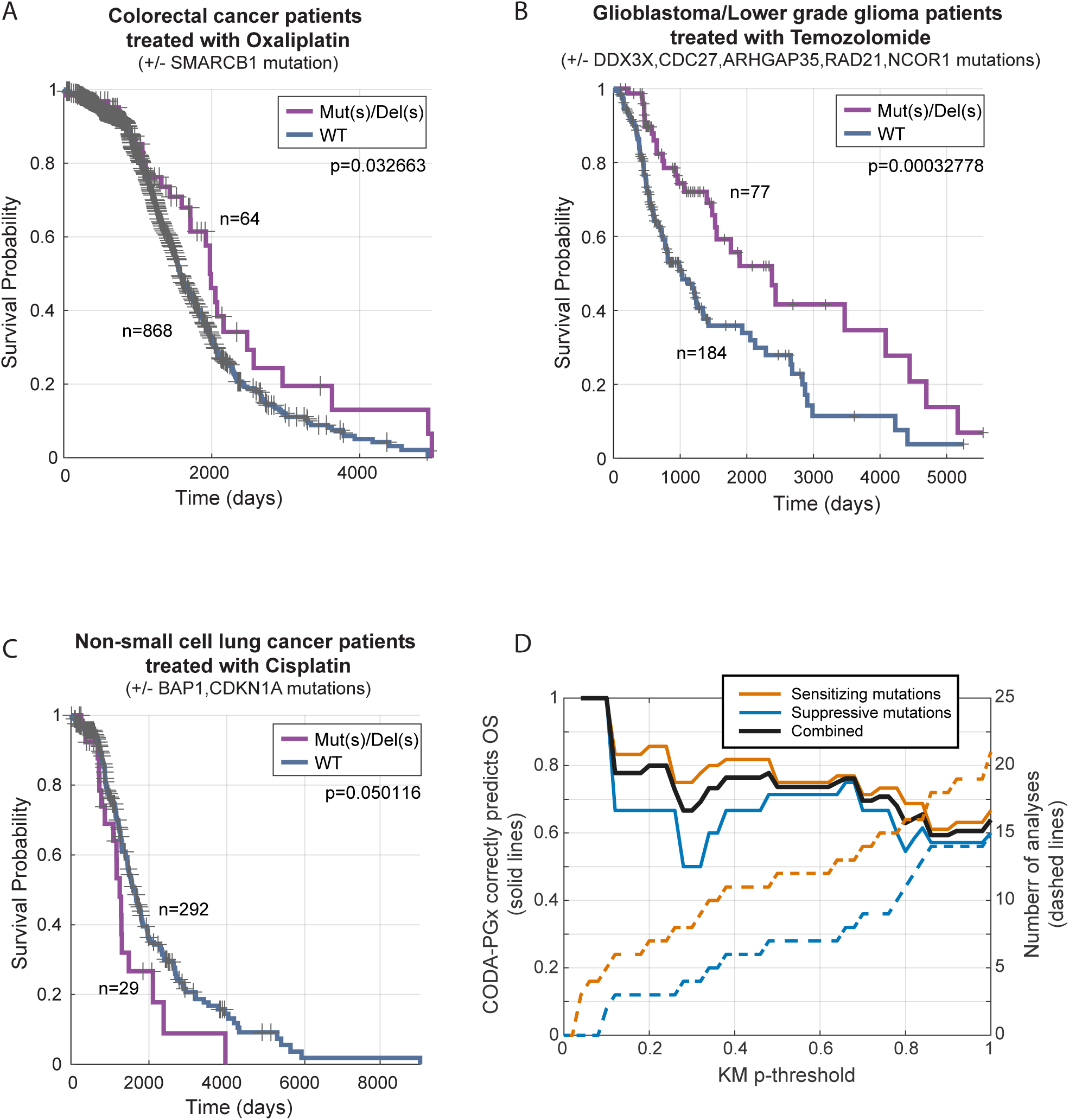
Retrospective clinical validation of CODA-PGX hits. **(A)** Plot of survival vs. time for colorectal cancer patients with (Mut/Del) mutations to SMARCB1 gene treated with Oxaliplatin compared to WT patients. **(B)** Plot of survival versus time for brain cancer patients with (Mut/Del) or without (WT) CODA-PGX-identified sensitizing mutations, treated with Temozolomide. **(C)** Plot of survival versus time for lung cancer patients with (Mut/Del) or without (WT) drug-suppresive mutations, treated with Cisplatin. **(D)**CODA-PGX predictions are generally associated with expected OS outcomes: the majority of sensitizing mutation/drug combinations are associated with improved OS, and the majority of suppressive mutation/drug combinations with worse OS.

These data suggest that the CODA-PGX platform can identify common cancer-causing mutations with potential to guide treatment decisions in the clinic. To more broadly assess whether CODA-PGX is predictive of clinical outcomes, we repeated the analysis above for all drugs and asked what proportion of the time CODA-PGX hit driver mutations predict the expected trend in OS compared to patients without these mutations in their tumors. Despite availability of RWE and molecular data from thousands of patients, subsetting on cancer type, the drug in question being used as frontline therapy, and presence of the mutation(s), yielded 41 drug-mutation combinations (i.e. “analyses”) where we were able to assess an association with overall survival (OS; **Figure 5D**). Both sensitizing and suppressive mutations yielded an enrichment for OS in the expected directions, with sensitizing mutations associated with improved OS, and vice versa. These effects were not caused by the mutations alone (**Supplemental Figure 6D**). Agreement was higher at more significant OS differences (i.e., to the left of **Figure 5D**), suggesting that statistical power may be limiting the overall analysis and that agreement overall would be higher if more patient data were available. Indeed, when requiring that at least 10 patients with the mutation exist in the analysis, 12/16 (75%) are associated with the predicted OS direction and the majority are at KM logrank p<0.5. Taken together, these data support that CODA-PGX predicted driver mutation – drug relationships are enriched for patient selection criteria with clinical relevance.

## DISCUSSION

To meet a significant unmet clinical need, we designed CODA-PGX, a platform to increase the discovery rate and translatability of novel targeted cancer therapies. We demonstrated that this approach delivers clinical trial-ready preclinical data that will enable rapid translation. We expect that CODA-PGX will provide an accelerated path to drug repurposing that will improve clinical benefits for patients and be transformative for cancer treatment. Specifically, we showed that CODA-PGX validates both established SL therapies and clinically using retrospective analyses. We further showed that CODA-PGX can identify novel SL interactions and open new avenues for targeted therapy. Our observation that platinum-based chemotherapies synergize with driver mutations similar to PARP inhibitors underlines the ability of CODA-PGX to identify new drugs to be used for targeted therapy. The implementation of these therapies may also allow these drugs to be used at lower doses in the patients with sensitizing driver mutations, which may result in a reduction of both toxic side effects in patients and cost to the healthcare system.

Although in this manuscript we focused on the SL interactions of PARP inhibitors and platinum-based chemotherapies, we expect that many additional insights into driver-drug interactions can be uncovered from further exploration of this dataset. For example, we note that MEK inhibitor Trametinib clusters with BRAF inhibitor Dabrafenib. Although this is expected, as BRAF acts upstream of MEK in the MAPK signalling pathway, this is a complex pathway with many levels of feedback regulation (*51*). Indeed, progress is being made to address the narrow therapeutic window of these inhibitors by developing allosteric inhibitors and testing combinations that may limit development of resistance (*52*). Further studies are needed to explore the mechanisms underlying these cancer driver-dependent dependencies to establish effective therapies.

While we demonstrated that CODA-PGX predictions are clinically relevant through retrospective analysis using publicly available cancer databases, we acknowledge that these analyses were limited by the number of patients for which data exists. Specifically, we tried to threshold on Varient Allele Frequency (VAF; fraction of reads in patient supporting mutation), maximum patient age, higher treatment regimens (e.g. 2^nd^/3^rd^ line treatment) but ran into power issues. We do, however, note that for well-powered drugs, higher VAF thresholds improved overall survival correlates.

Classical pooled CRISPR/Cas9-based screens have been widely applied to identify SL interactions between genetic mutations and chemotherapeutic agents (*8, 22, 53–55*). However, these approaches explore one condition versus all possible mutations. They identify huge numbers of possible interactions but lack the ability to prioritize which are actionable. In contrast, our focus on driver biology in CODA-PGX is unique in that it guarantees a causal SL relationship between drivers and screening hits in the presence of a given drug. However, we note that the driver perturbation itself may not reflect the initial biological wiring in a tumor/clinical setting because tumors have evolved from their initial driver events. It is for this reason that we believe that the intersection of consortia efforts and focused isogenic SL screening is a powerful filter for identifying genuinely causal and clinically-preserved SL vulnerabilities, thereby yielding the most therapeutically valuable targets. We expect that CODA-PGX will reveal novel drug repurposing opportunities that can be rapidly validated, thereby enabling a fundamental shift in the development and translation of cancer therapy, increasing patient quality of life and prolonging survival.

### Limitations of the study

We acknowledge the following limitations of the CODA-PGX method. First, CODA-PGX screening is not comprehensive: it is impossible to know if we are evaluating all possible driver genes in any given screen. Despite this, we believe that the drug-driver relationships that CODA-PGX does evaluate will be transformative for clinical treatment of a wide variety of cancers. Second, our ability to perform retrospective analysis is currently limited by the size of patient databases. It is expected that these databases will continue to expand and as they do, the implications of the CODA-PGX platform analysis will be fully realized.

## MATERIALS AND METHODS

### Study design

The overall objective of this study was to develop a pharmacogenomic screening platform that would accurately identify SL drug-gene mutation combinations which will translate into new targeted cancer therapies. To this end we generated a pooled CRISPR guide KO library that targeted driver genes frequently mutated in cancer. Proof-of-concept library screens were conducted and optimized using 4 different PARP inhibitors. Optimized conditions, including drug dose and length of drug treatment, were applied to screen 20 anti-cancer drugs used in cancer clinics. To ensure accuracy, screens were conducted in multiple human cell lines that represented a range of common cancer types. SL relationships identified in these screens were confirmed by generating isogenic cell lines and comparing the growth of Parental and KO cells over a range of drug doses. Both CDX and PDX models were used to further validate in vitro SL findings. Female mice were used in both models. Studies were performed according to protocols approved by the Institute’s Animal Care Committee. A retrospective analysis of clinical outcome datasets allowed us to compare our platform predictions to actual overall survival of cancer patients.

### Plasmids

plentiCRISPR v2 was a gift from Feng Zhang (Addgene plasmid # 52961; http://n2t.net/addgene:52961; RRID:Addgene_52961). psPAX2 was a gift from Didier Trono (Addgene plasmid # 12260; http://n2t.net/addgene:12260; RRID:Addgene_12260). pMD2.G was a gift from Didier Trono (Addgene plasmid # 12259; http://n2t.net/addgene:12259; RRID:Addgene_12259).

### Cell lines

MDA-MB-231, A549, LS180 and OvCar-3 cell lines were purchased from the American Type Culture Collection. KP-4 cells were purchased from Japanese Collection of Research Bioresources Cell Bank. Cell lines were maintained in DMEM with 10% FBS and 1% antibiotics/antimycotics, except OvCar-3 cells which were grown in 20% FBS supplemented with 10 ug/ml Insulin (Sigma). All cell lines tested negative for Mycoplasma and were authenticated by Short Tandem Repeat analysis.

### Pooled sgRNA library design and cloning

Our CRISPR knockout pooled sgRNA lentiviral library targets 180 known cancer driver genes (**Supplemental Table 1**). The library was designed based on the published sgRNA sequences in the optimized human CRISPR knockout pooled Brunello library (*56*) that contains 4 sgRNAs per gene. Forward and reverse DNA oligonucleotides were synthesized (Eurofins Genomics) for each sgRNA according to the following sequence: Forward 5’-***CACC***G(N20)-3’, Reverse 5’-***AAAC***(N20)C-3’, including overhang sequences for cloning (bold italics). Following a general CRISPR guide cloning protocol (https://www.addgene.org/crispr/zhang/), each pair of oligonucleotides was combined, phosphorylated, denatured and annealed. Equal amounts of all annealed guides were pooled together, diluted and cloned into BsmBI-digested (NEB)/ CIP-treated (NEB) plentiCRISPR v2 (Addgene) using Blunt/TA Ligase (NEB). The library was amplified by large-scale transformation of Stbl3 cells (ThermoFisher Scientific) and grown on LB plates containing ampicillin. Colonies were scraped off the plates and plasmids were extracted with a plasmid maxiprep kit (Qiagen).

### Lentivirus generation and target cell transduction

Lentivirus was produced according to the “General Protocol for Generation of lentivirus in 293T cells” using “PolyJet™ In Vitro DNA Transfection Reagent” (Signagen Laboratories). Briefly, 293T cells were grown in 100 mm dishes until approximately 90% confluent and transfected with equimolar amounts of pLentiCRISPRV2/guide library plasmid, psPAX2 (Addgene) and pMD2.G (Addgene) plasmids. Medium containing lentiviral particles was collected at 48 and 72 hours post transfection and concentrated using PEG-it Virus Precipitation Solution (System BioSciences). Precipitated lentiviral particles were resuspended in DMEM, aliquoted into 50-100 ul volumes and frozen at −80°C. For each target cell line, the lentiviral titer was determined by transducing cells with serial dilutions of lentivirus, followed by puromycin treatment and counting surviving cells. To generate transduced cells for the drug screens, cells were transduced at 0.3 x multiplicity of infection (MOI). After 24-48 hours, puromycin (6 ug/ml) was added to the medium and the cells were selected for 5 to 7 days. Transduced cells were counted, cryopreserved and stored in LN2 or used in the drug screen.

### Drug screening

For each drug/cell line combination we generated a viability/dose response curve (DRC), which enabled us to calculate drug concentrations that corresponded to different IC values (**Supplemental Table 3**). Interrogation of PARP inhibitors across IC5-IC50 revealed best responses in the IC5-20 range, regardless of cell line background (**Supplemental Figure 4**). For each set of drug screens, a Day 0 cell pellet of approximately 500K cells was saved for analysis. To conduct the screen, 300K cells were initially plated in 60 mm dishes. After approximately 24 hours the culture medium was replaced with fresh medium containing the appropriate drug. Drug and culture medium were replaced every 48 hours. At 100% confluency the number of cells was recorded, and 800K cells were replated into a 100 mm dish and the drug treatment was continued until 100% confluency. The cells were counted again and saved for NGS analysis. For each cell count, the population doubling was determined (log10 (N/No) × 3.32, where N = final number of cells and No = starting number of cells) and then added together to determine the total number of doublings. Experiments were performed in duplicate as biological replicates.

### NGS library generation and sequencing

For each sample a minimum of 300K cells was analyzed. Genomic DNA (gDNA) was extracted with a DNeasy Blood and Tissue kit (Qiagen). A two-step amplification protocol adapted from Chan et al. (*53*) was used to generate NGS libraries. In brief, the protocol was followed except: 14 ug (3.5 ug x 4 reactions) of gDNA were used for PCR1, using in-house forward (5’-TCATATGCTTACCGTAACTTGAAA-3’) and reverse (5-’AACTTCTCGGGGACTGTGG-3’) primers. The number of cycles was extended to 8 and 28 for PCR1 and PCR2, respectively. PCR products were purified with HighPrep PCR Clean-up Beads (MagBio Genomics) and quantified via Nanodrop spectrophotometer (Thermo scientific). The sequencing library was generated by combining equal amounts of purified PCR products and quantified using the NEBNext Library Quant Kit for Illumina (New England Biolabs). The primer 5’-ACACTCTTTCCCTACACGACGCTCTTCCGATCTTTGTGGAAAGGACGAAACACCG-3’ was used to sequence on Illumina platforms.

### Screening data analysis

Demultiplexed (per sample) fastq files were aligned against guide library sequences (**Supplemental Table 2**) using blat, allowing up to 2 mismatches per read. Driver gene guide sequences were designed using previous screens (*56*). 96 negative control guides were designed using ChopChop (*57*) against intergenic non-conserved (mean PhastCons (*58*) score less than 0,1) regions of the genome. 85-92% of reads aligned uniquely to guide sequences across all the screens. Counts per guide were normalized to counts per million (CPM) by dividing by the total number of aligned reads for that sample, in millions. CPM-level was quantile-normalized across samples and analyzed by PCA to ensure that data was free of systematic biases (e.g. batch effects arising from screening or NGS library generation). Normalized guide-level data for each drug was converted to ratios by dividing by the average of DMSO/vehicle control data generated in parallel. All screening batches had a minimum of 6 DMSO/vehicle control screens included. Gene-level fold-change was computed as the average log-base-2 of the 3-4 guide ratios, and gene-level significance was computed using a ranksum test of the 3-4 guide rations vs the 96 negative control ratios. Significance therefore reflects a consistent shift in the 3-4 gene-level guides against the null (negative control) distribution that is measured and processed identically. When the cell line contained a known oncogene mutation (*25*), the negative control and gene-specific signals were reversed (i.e. mutation present in negative controls, but absent in gene-specific KO). False-discovery rates were gauged against a distribution of significance values generated from all possible vehicle versus vehicle controls (i.e. pretending each DMSO control is a drug-treated sample, and comparing against the 5+ other DMSO controls).

### Principal Component Analysis (PCA)

Normalized and filtered gene-level (see above) fold-change values were subject to PCA analysis using the linear model of variance in R, and visualized using ggplot2.

### Cell line knockouts

Generation of STAG2 KO OvCar-3 cells was accomplished using Synthego multi-sgRNAs and lipofection-based transfection of RNPs (ribonucleoprotein complexes). Multi-sgRNAs tiling the first exon of STAG2 were purchased from and designed by Synthego (Gene Knockout Kit v2, chemically modified to prevent its degradation). sgRNA sequences are as follows: ACUUGUAAAAAAGGCAAAAA; UCUGGUCCAAACCGAAUGAA; UUGUUUGAAGUUGUUAAAAU. Cas9 2NLS nuclease (S. pyogenes protein) was also purchased through Synthego. As per Synthego’s Immortalized Cell Lipofection protocol, RNPs were complexed at a sgRNA:Cas9 ratio of 1.3:1. RNPs were delivered via the Lipofectamine CRISPRMAX Transfection Reagent to 50,000-100,000 cells in a 1.5mL microcentrifuge tube. Cells were subsequently seeded into a 24 well plate, grown to confluency and then expanded into a 60 mm dish. To generate sufficient KO cells for the CDX studies, a large-scale transfection of 4M cells was performed where the RNP reaction and transfection were scaled up by a factor of 40. Transfected cells were grown for approximately 7 days until the cells had expanded to 20M. The STAG2 KO efficiency was determined to be 100% immediately prior to the CDX study. The KO efficiency was determined using the Synthego Performance Analysis, ICE Analysis. 2019. V3.0. Synthego, following amplicon amplification using forward primer – GGACACCACAAAGAGGCTGT; Reverse primer – AATACATCCCAAGAGTTTTCTGATG.

### Dose/viability response curves (DRCs)

Cell viability following drug treatment was assessed using the Promega CellTiterGlo 2.0 viability assay, a luminescence-based read out of ATP (generated by metabolically active cells). Cells were seeded at 500-2000 cells per well into 96 well white luminescence assay plates. Following attachment, cells were treated with seven drug concentrations ranging from 50 µM to 0.0032 µM in a five-fold dilution gradient. To achieve a final vol/vol DMSO concentration of 0.5%, drugs were initially diluted to 200x in DMSO, then diluted further into medium. In addition to drug-treated wells, control wells of cells treated with 0.5% DMSO and wells containing media only (no cells) were included in the plate. Each data point was performed, at a minimum, in triplicate. Cell viability was read on the SpectraMax Gemini XPS when DMSO control wells reached approximately 70% confluency. Luminescence measurements were background subtracted by the average signal of media-only wells and further normalized to the average signal of DMSO-treated wells.

### Cell-derived xenograft (CDX) experiments

NOD.Cg-PrkdcscidIl2rgtm1Wjl/SzJ (NSG) mice were housed and handled in accordance with the approved guidelines of the Canadian Council on Animal Care. All experiments were approved under the Animal Use Protocol of the Research Institute of McGill University Health Centre (AUP#7802). Housing condition for mice: Temperature = 21°C +/- 1°C; Humidity = 40– 60% RH +/- 5% RH; Lighting = 12 hr ON / 12 hr OFF daily cycle. WT and STAG2 KO OvCar-3 cells were cultured in DMEM with 20% FBS and 10 μg/mL insulin for 6 days at 37°C in 5% CO2. OvCar-3 cell line was tested negative for Mycoplasma by Diagnostic Laboratory from Comparative Medicine and Animal Resources Centre before in vivo injection. Cells were diluted in the 1:1 PBS/Matrigel mixture, and 3 million cells per mouse were subcutaneously injected into the right flank of 12-week female NSG mice. Tumor volumes were measured with a digital electronic caliper three times per week once tumors are palpable. Once the average tumor volumes of both WT and KO groups reached approximately 200 mm^3^, all the mice from the two groups were randomized into vehicle (saline) and 40 mg/kg Carboplatin (MedChemExpress) treatments. Vehicle and Carboplatin were delivered by intraperitoneal Injection every 4 days. Tumor volume was calculated according to the following formula: [4/3 × π × (length/2) × (width/2)2] to generate a growth curve. Tumor Growth Inhibition (TGI) is defined as (1 – (mean volume of Carboplatin treated tumors)/(mean volume of vehicle tumors)) × 100%.

### Patient-derived xenograft (PDX) experiments

Carboplatin standard of care (SOC) PDX data was generated by Crown Bio and is being used with permission. It was identified using HuBase and was generated using Crown Bio standard operating protocols (*59*).

### Retrospective clinical outcome analysis

Project GENIE NSCLC and CRC mutation and clinical outcome datasets were downloaded from Synapse (GENIE ID: syn7222066, BPC dataset IDs: syn30991602, syn27056179). LoF variants included: nonsynonymous amino acid changing variants with a predicted functional impact on protein using MutationTaster (*60*), nonsense inducing variants, and single allele or double-allele deletions (i.e. present as zero on single-copy). GoF variants were considered to be hotspot nonsynonymous substitutions (*61*) in known oncogenes. WT calls were made when no nonsynonymous variants were present (including unfiltered variants) and >95% of all exons of that gene was represented on the sequencing platform used to generate the data (GENIE has 104 different platforms, mostly Illumina-based but differing in genes being captured). TCGA data was downloaded from Genomic Data Commons (*62*) and processed the same as above. For each drug screen (i.e. row in **Figure 3A**), mutated gene sets were extracted as meeting the following criteria: ranksum gene-level p<0.05, and gene-level fold-change is less that −2 for sensitizing mutations, and greater than 2 for suppressive mutations. Sensitizing and suppressive gene sets were treated separately. Patients in the cancer type with the highest number of patients treated with the drug in question were identified (from TCGA and BPC) and split into two groups: patients with one or more of the hit mutations (using criteria above), and patients WT for the mutations. Kaplan-Meier logrank test was conducted to test the null hypothesis that there is no difference in OS (*63*). Patients without a known OS outcome at the final time of reporting were included and censored (shown as + marks in **Figure 5**).

### Statistical analysis

In CDX studies, all results are presented as the mean±SEM. The difference between vehicle and treatment group was analyzed by two-way ANOVA using GraphPad. Kaplan-Meier survival curves were analyzed by logrank test. \**P* < 0.05 was considered statistically significant.

## List of Supplementary Materials

Fig S1 Number of small molecule drugs that were discontinued after entering clinical trials.

Fig S2 Treemap depicting the genetically targeted treatment landscape of cancer.

Fig S3 DepMap cell fitness of cells after knockout of the gene.

Fig S4 Four approved PARP inhibitors screened in MDA-MB-231.\

Fig S5 Carboplatin screens in KP4 and MDA-MB-231 cells.

Fig S6 Kaplan-Meier Survival Curves.

Table S1 (ST1) Driver genes screened in this study and evidence supporting their driver roles.

Table S2 (ST2) Guides and oligo sequences used in this study. Brunello guides were published previously (PMID: 26780180) and negative controls were designed using ChopChop.

Table S3 (ST3) Data underlying Figure 3, along with unfiltered p-values and log(fold-change) values; drug concentrations and ordering catalog numbers used are also included.

Table S4 (ST4) Literature support for screening discoveries shown in Figure 3.

## ACKNOWLEDGMENTS

We thank Greg Vontz, Joe Bondy-Denomy, and Brian DeVeale for critical review of the manuscript.

## AUTHOR CONTRIBUTIONS

Conceptualization: TB

Methodology: PT, JC, DCV, MD, YZ, RM, SY, TB

Investigation: PT, JC, MD, TB

Funding acquisition: AWC, TB

Project administration: PT, JC, MD, AWC, TB

Supervision: PT, JC, MD, AWC, TB

Writing – original draft: JC, SH, TB

Writing – review & editing: PT, MD, SH, AWC, TB

---

**Supplemental Figure 1.**
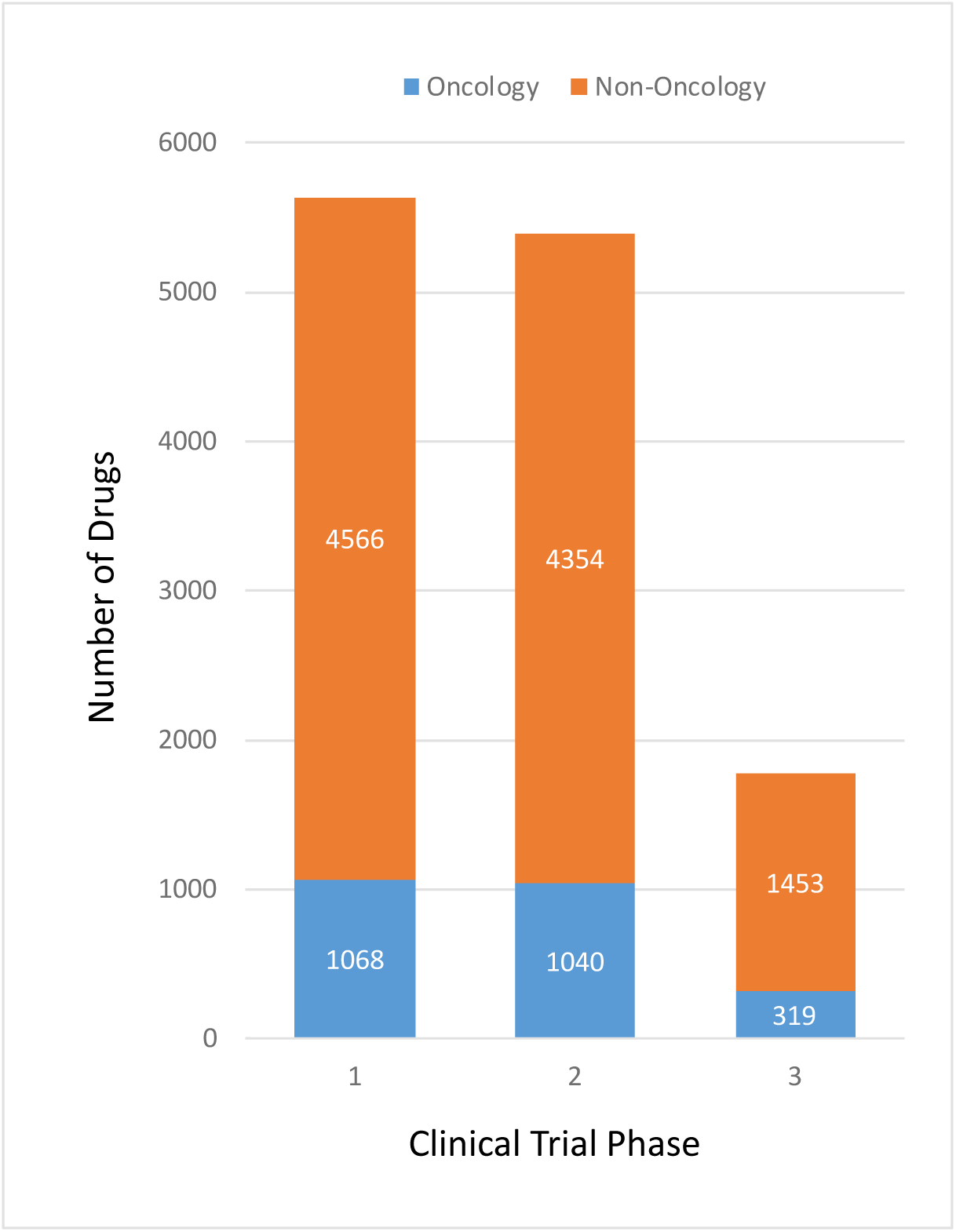
Number of small molecule drugs that were discontinued after entering clinical trials, split according to the maximum phase they were tested in. Of 12,800 total discontinued drugs, 2,427 were used to treat oncology indications.

**Supplemental Figure 2.**
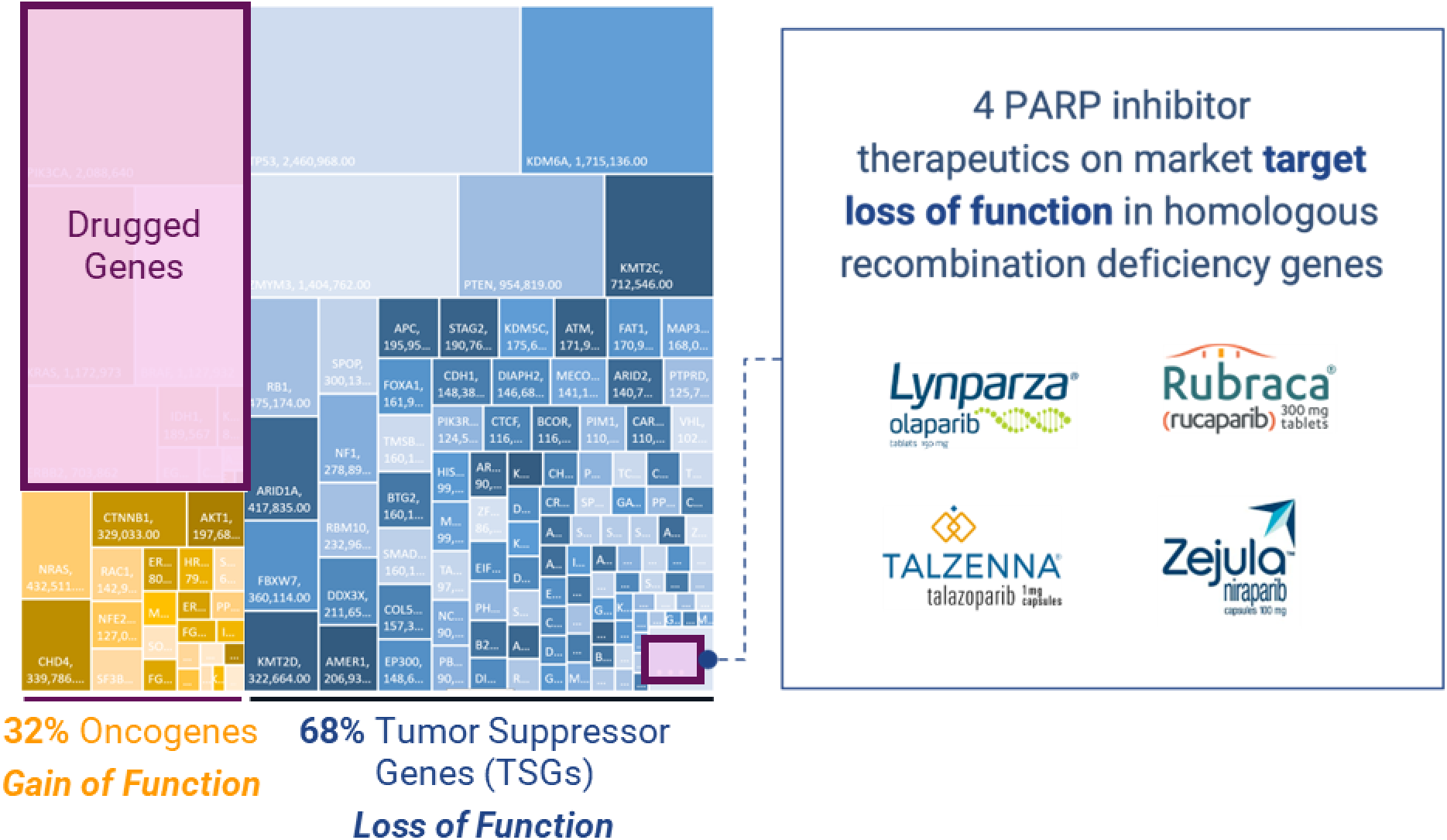
Treemap depicting the genetically targeted treatment landscape of cancer. The mutation frequencies (from TCGA and GENIE) within each cancer tupy of 184 of the most common driver genes (see **Supp Table 1**) were multiplied by the prevalence (number of patients; taken from ACS and NCI) of each cancer type in the United States, and added together. In other words, if every driver gene had an available targeted therapy, this treemap would depict the number of patients that each therapy could be used to treat. There are approximately 15 million patients living with cancer in the United States, and the numbers above add up to ~25 million, since most patients have more than one driver mutation. The vast majority of currently available cancer targeted therapies on the market treat oncogenic mutations.

**Supplemental Figure 3.**
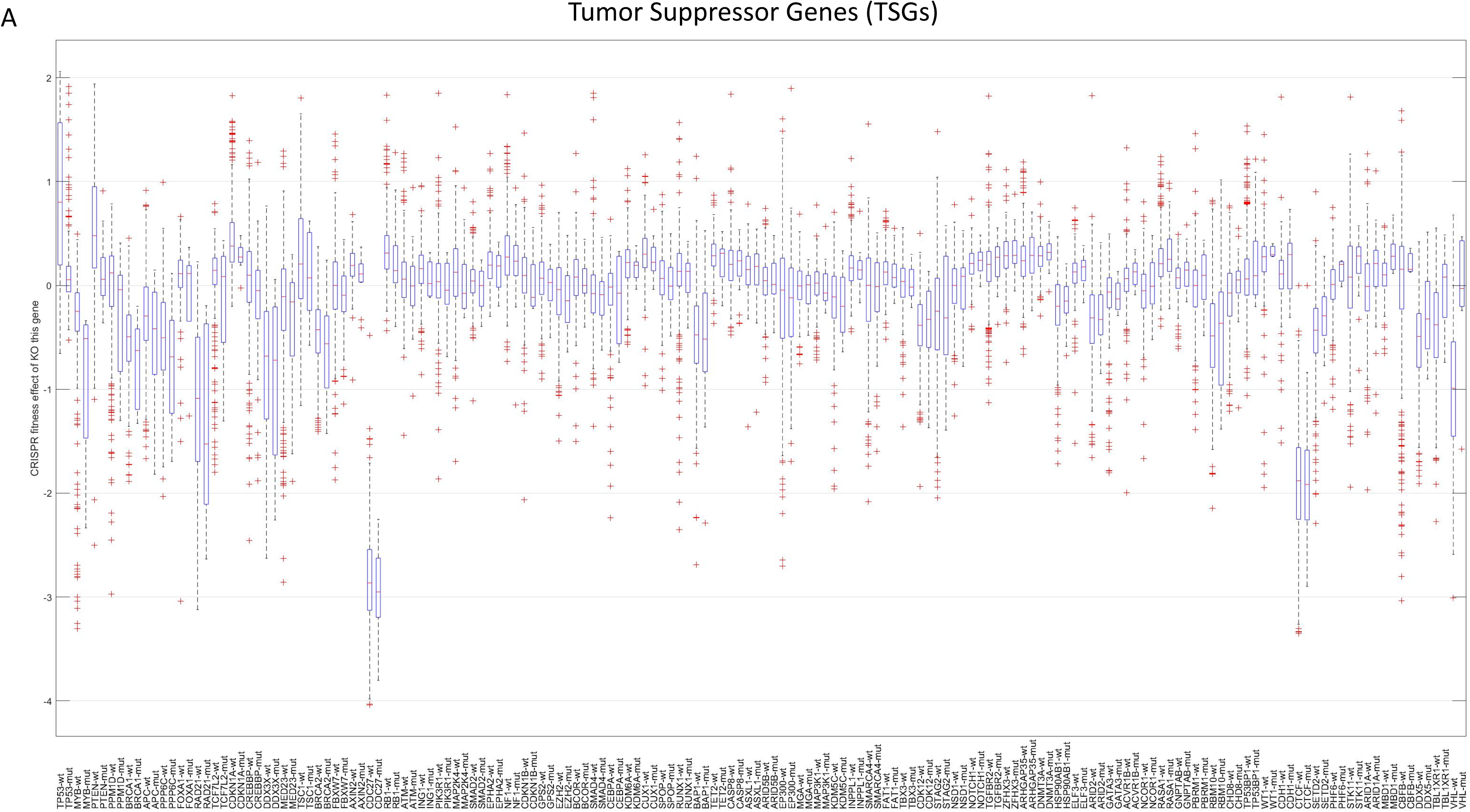
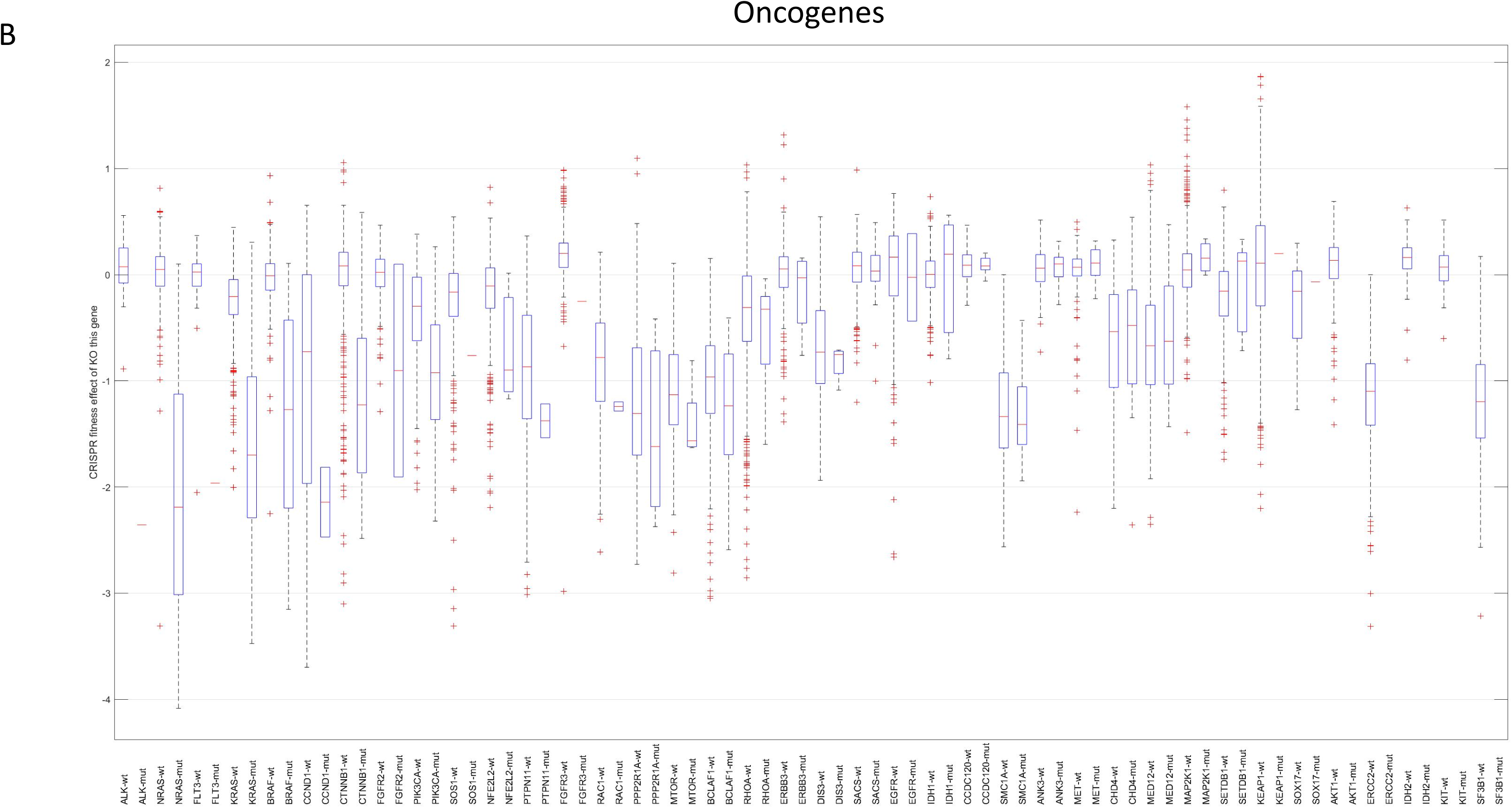
DepMap cell fitness of cells after knockout of the gene on x-axis, with cells split by mutant vs wt for that gene. **A**) Tumor Suppressor Genes (TSGs): in general KO of wt TSG leads to an increase in fitness (proliferation) while mutant remains near 0 (i.e. unchanged), like ly because it already has a loss of function in that gene. **B**) Oncogenes: knockout of oncogenes with gain of function mutations, tends to be deleterious to cells relative to the knockout of wt.

**Supplemental Figure 4.**
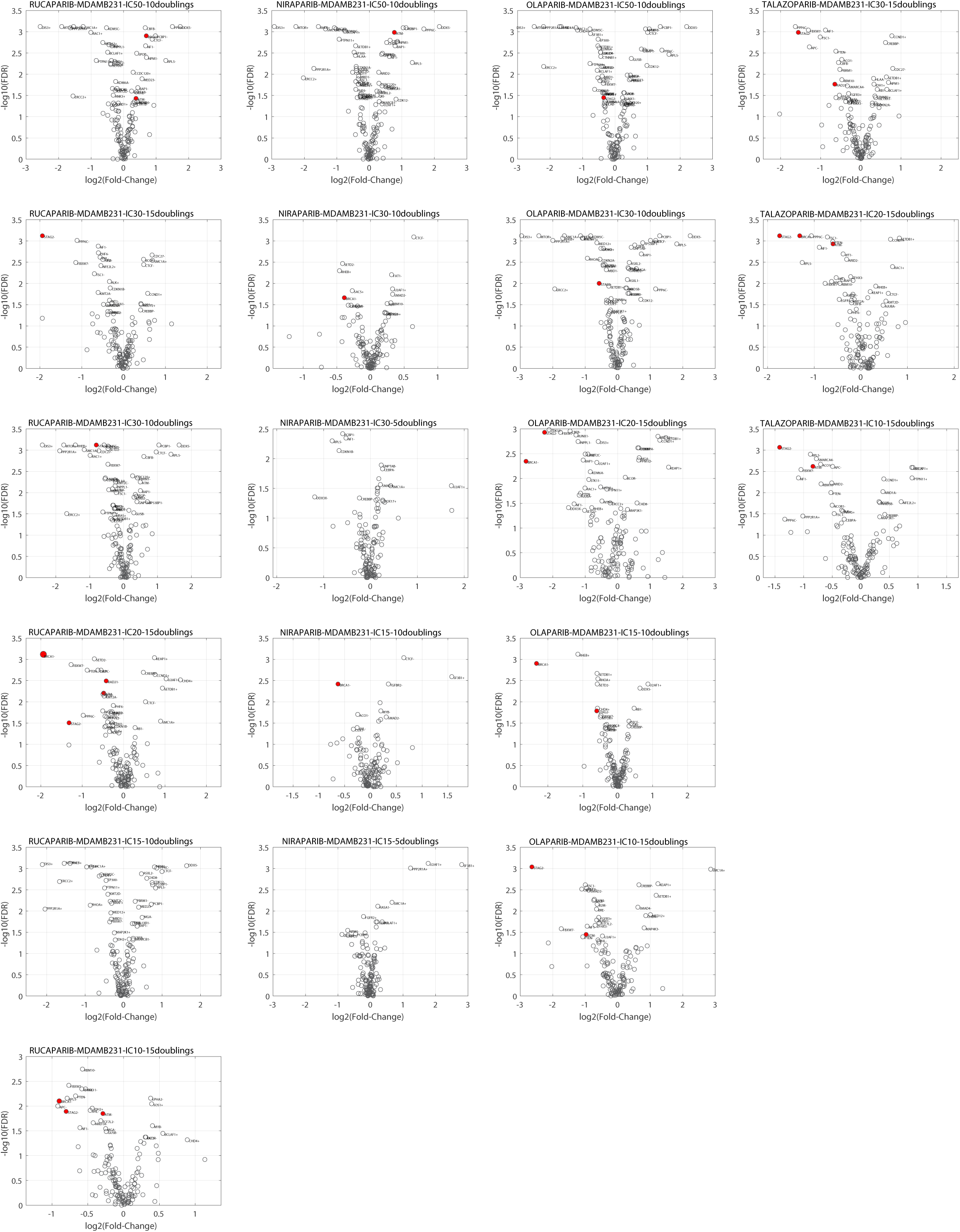
Four apprvoed PARP inhibitors screened in MDAMB231 (breast) cells across a range of inhibitory concentrations (IC5-IC50) for either 10 or 15 population doublings. IC10/IC15/IC20 and 10 population doublings consistently yielded the expected sensitizing mutations (BRCA1, ATM, STAG2, RAD21 - shown in red).

**Supplemental Figure 5.**
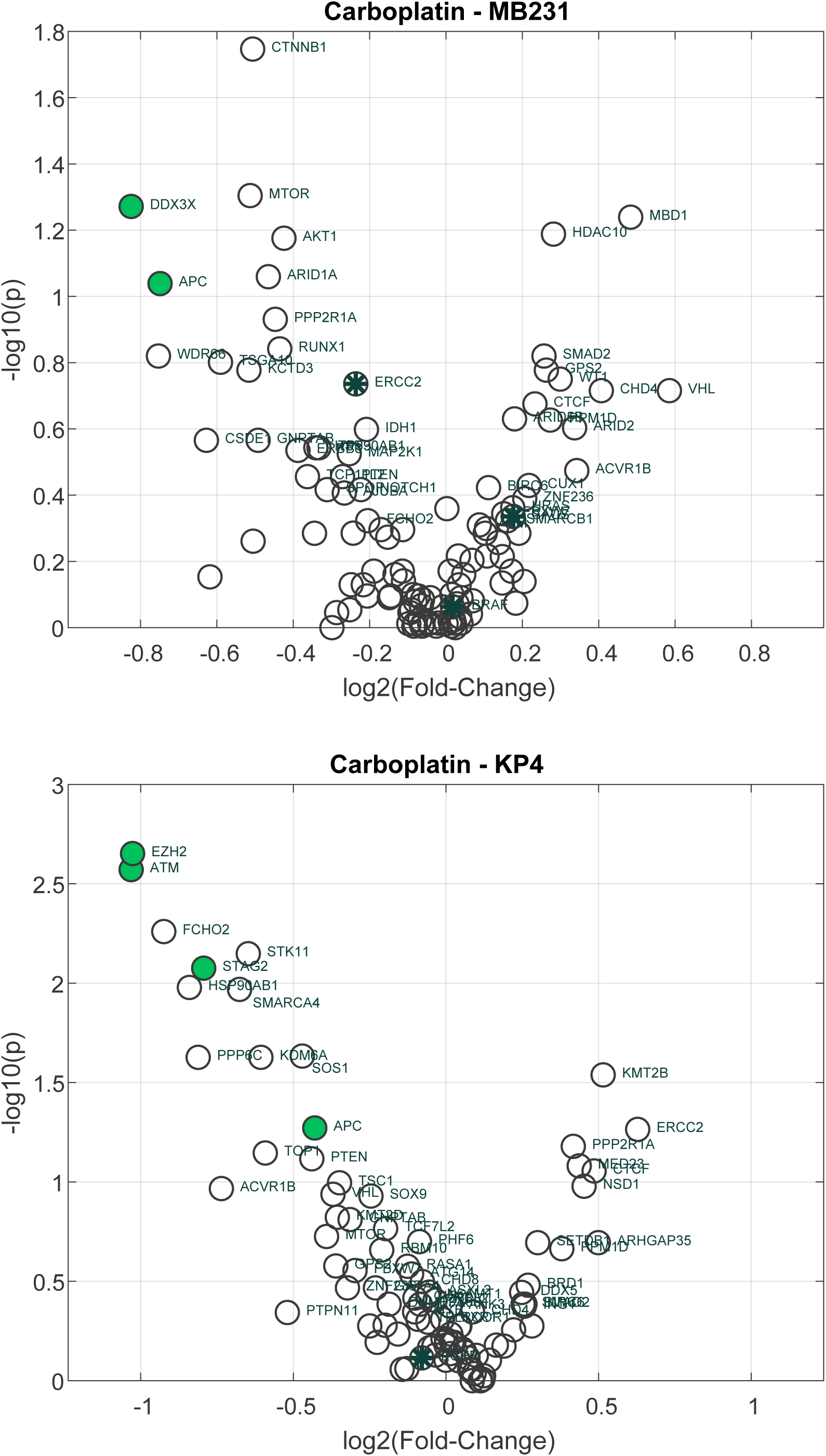
Carboplatin screens in KP4 and MDA-MB231 cells.

**Supplemental Figure 6.**
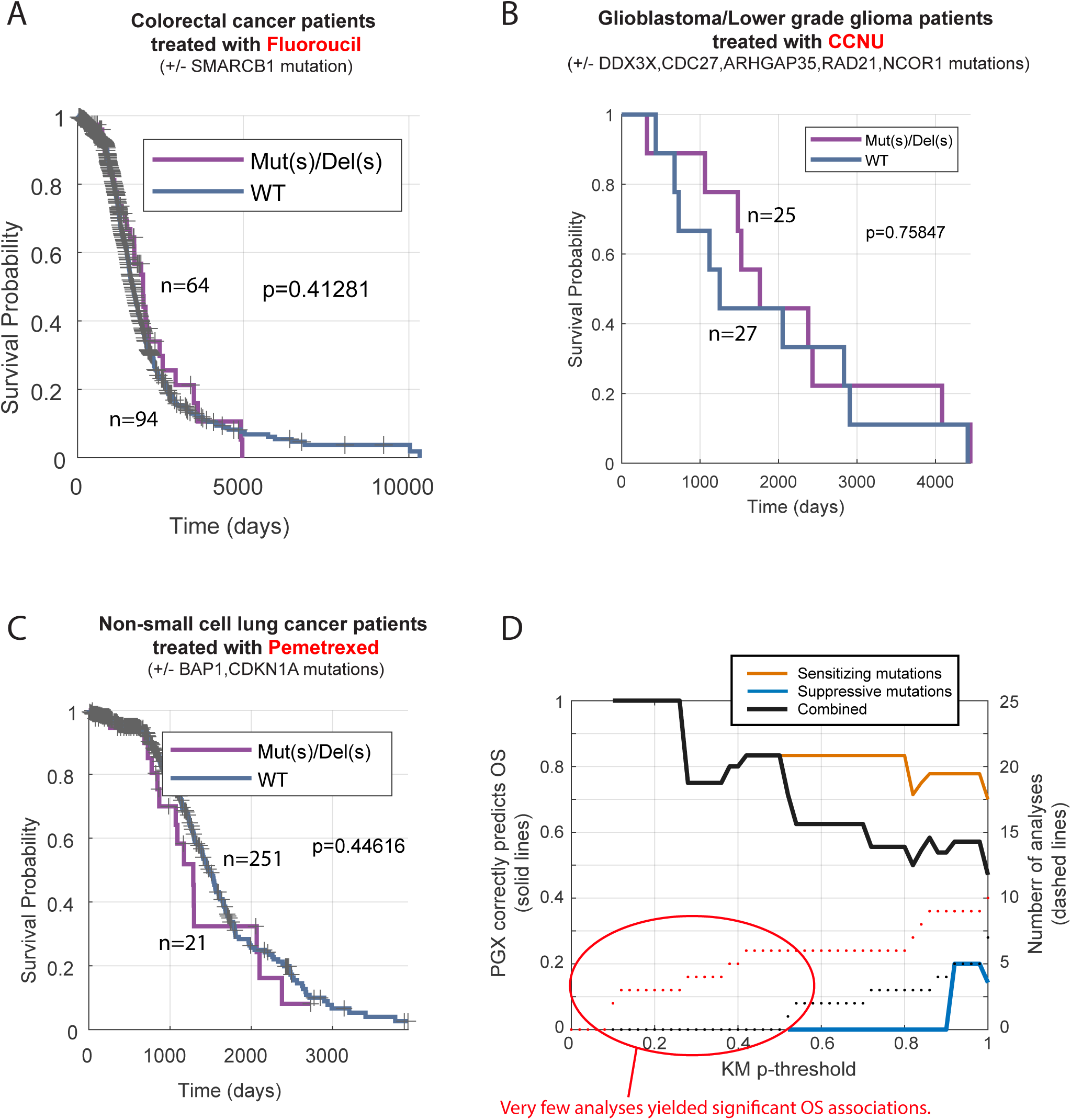
Identical analysis as shown in Figure 5, but using another well represented drug available in the RWE databases.

